# Balancing conservation priorities for grassland and forest specialist bird communities in agriculturally dominated landscapes

**DOI:** 10.1101/2021.08.05.455200

**Authors:** Devin R. de Zwaan, Niloofar Alavi, Greg W. Mitchell, David R. Lapen, Jason Duffe, Scott Wilson

**Author notes:** LRH: *de Zwaan et al*. RH: Conserving species-at-risk in farmland.

## Abstract

Effective conservation planning often requires difficult decisions when at-risk species inhabit economically valuable landscapes or if the needs of multiple threatened species do not align. In the agriculture-dominated landscape of eastern Ontario and southwestern Quebec, Canada, conflicting habitat requirements exist between threatened grassland birds benefiting from certain agriculture practices and those of a diverse woodland bird community dependent on forest recovery. Using multi-scale species distribution models with Breeding Bird Survey (BBS) data, we assessed habitat suitability for 8 threatened grassland and forest specialists within this region. We also identified landscapes that jointly maximize occurrence of the 8 focal species and diversity of the overall grassland and forest communities. Influential habitat associations differed among species at the territory (200m radius) and landscape level (1km), highlighting the importance of considering multiple spatial scales. Species diversity was maximized when forest or grassland/pasture cover approached 40–50%, indicating a positive response to land cover heterogeneity. We identified species diversity hotspots near Lake Huron, as well as along the shore and southeast of the St. Lawrence River. These areas represent mosaic landscapes, balancing forest patches, wetland, grassland/pasture, and row crops such as corn, soybean, and cereals. Despite drastic landscape changes associated with agroecosystems, we demonstrate that targeted habitat protection and enhancement that prioritizes land cover diversity can maximize protection of bird communities with directly contrasting needs. We highlight multiple pathways to achieve this balance, including forest retention or separating row crops with hedgerows and wooded fence-lines, improving flexibility in conservation approaches.

## 1. Introduction

Conservation planning requires decisions on how to allocate limited resources to maximize outcomes (Margules and Pressey 2000, Naidoo and Ricketts 2006). For institutions that have multiple conservation goals under their mandate, these decisions may require trade-offs among goals that do not align, either because of their location or the strategy needed. For example, a poor alignment of goals could occur when proactive versus reactive conservation strategies emphasize different geographic areas (Brooks et al. 2006, Wilson et al. 2019) or where greater investment in the recovery of a small number of highly threatened species comes at the cost of protecting a larger number of less threatened species (Martin et al. 2018). In this study, we demonstrate how we can prioritize agricultural landscapes for conservation in a historically forested ecosystem. In this case, there are conflicting goals between the needs of threatened grassland species that benefit from certain agricultural practices and those of a diverse community of woodland-associated species that benefit from forest recovery and retention.

Prior to European settlement, eastern North American ecosystems were largely forested with small openings associated with wetlands, grassland, and Indigenous agriculture (Williams 1989, Butt et al. 2005). Following settlement, forests were cleared for crop agriculture and silviculture with expansion beginning in the east and continuing westward (Ramankutty and Foley 1999, Steyaert and Knox 2008). As the center of agricultural productivity shifted towards the mid-continent, agricultural areas in eastern North America were increasingly abandoned, promoting forest recovery in the late 19^th^ and 20^th^ centuries at rates that varied regionally (Pimm and Askins 1995, Ramankutty and Foley 1999, Ramankutty et al. 2010). In recent decades however, forest recovery has slowed or even reversed in eastern North America due to urbanization, agricultural intensification, and timber harvesting (Drummond and Loveland 2010)

The opening of the landscape in the eastern hardwood forests of North America would have benefitted species that inhabit grasslands, many of which were also losing extensive areas of habitat to agriculture in central North America following the near disappearance of tall grass prairie (Vickery et al. 1999, Norment 2002). Today, declining grassland species such as bobolink (*Dolichonyx oryzivorus*) and Eastern meadowlark (*Sturnella magna*) readily use agricultural landscapes in eastern North America, particularly those associated with pastoral or hay production (Wilson et al. 2017). These two species are federally listed under Canada’s Species at-Risk Act (SARA 2002) and thus there is a mandate to enact measures for their recovery. At the same time, crop and pastoral agriculture are associated with the loss of forest bird diversity with declines due mainly to the disappearance of insectivorous Neotropical migrants from the community (Endenburg et al. 2019). Some of these Neotropical migrants are also federally listed under SARA (e.g., wood thrush *Hylocichla mustelina*, Eastern wood-pewee *Contopus virens*).

This scenario leads to a challenging conservation dilemma on how to best manage agroecosystems when the habitat needs for declining, federally listed grassland species conflict with those for threatened forest birds and the broader diversity of the forest bird community.

In this study, we focused on an extensive, agricultural-dominated region of eastern Ontario and southwestern Quebec (Figure 1A) to address this conservation challenge.

**Figure 1.**
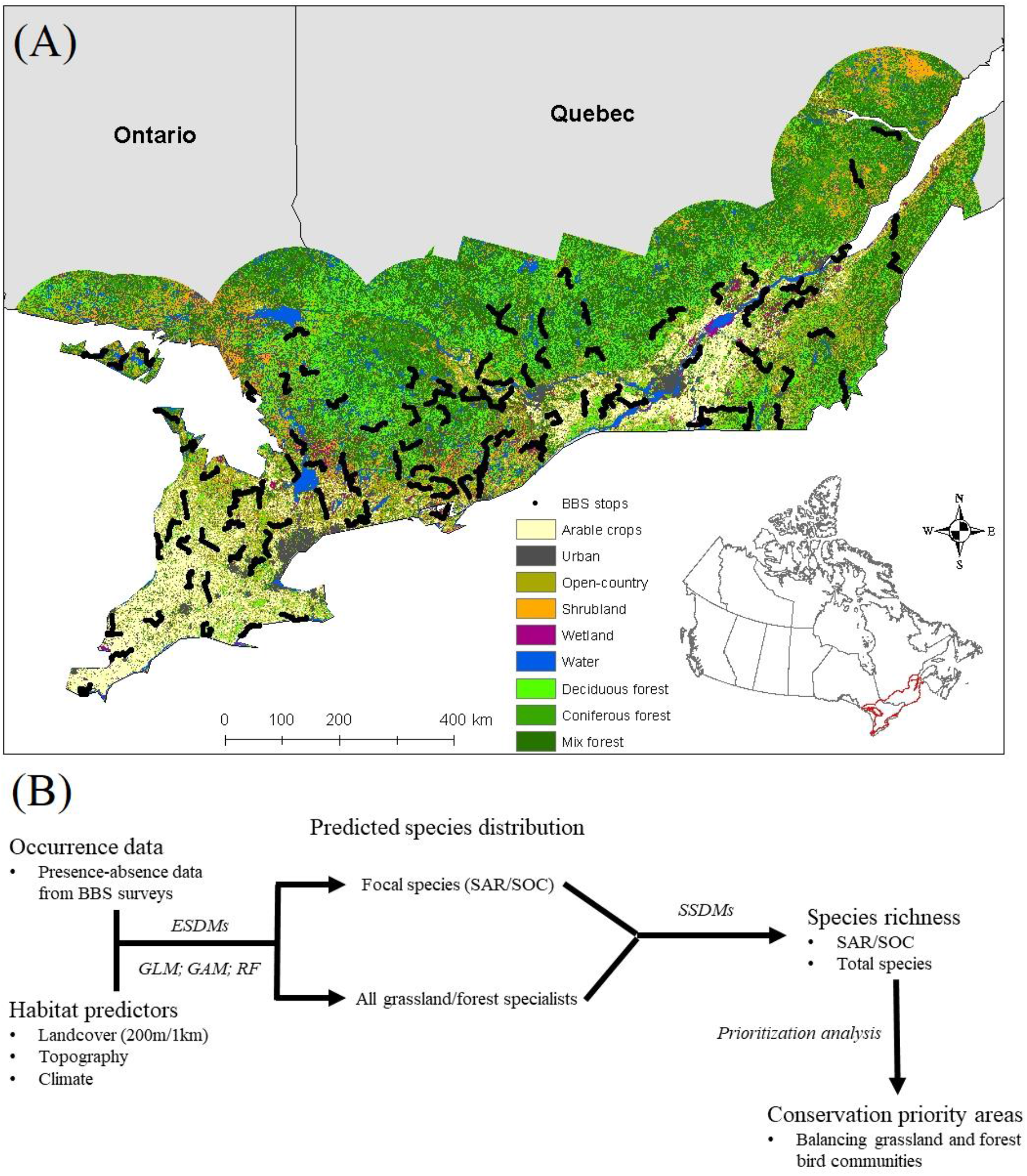
(A) Map of the study region showing the distribution of the main land cover variables and the individual Breeding Bird Survey (BBS) stop locations (black points). (B) A schematic of the modelling process.

Specifically, we (1) used a multi-scale species distribution approach to identify geographic areas with the greatest potential to maximize conservation of three targets: i) declining grassland birds, ii) declining forest birds, and iii) total avian diversity. For both declining grassland and forest birds we considered species that are federally listed in Canada as species at risk (SAR), but also species of conservation concern that have yet to be listed. We then assessed (2) which landscape characteristics and spatial scales most influenced occurrence of each species, and thus how to optimize conservation of the three target groups. Finally, (3) we used this information as the basis for a systematic prioritization analysis to formally identify the landscapes and regions that allow us to reach a 25% landscape area protection goal in a manner that maximizes conservation of grassland and forest-dependent species. This goal was chosen to align with Canada’s commitment to protect 25% of terrestrial area by 2025. The results from this study will provide critical insights into the types of landscapes we need to manage or protect to jointly benefit at-risk forest and grassland specialists despite contrasting habitat needs.

### 2. Material and Methods

### 2.1. Study region and species

The study region in eastern Ontario and southwestern Quebec, Canada, represents a land cover gradient from forest to intensive agriculture (primarily livestock cash crop) that spans approximately 600 km in latitude and 1000 km in longitude (Figure 1A). This region has experienced extensive habitat modification due to agricultural practices and urbanization, particularly in the south. To the north and west, forests have been cleared for timber and pastoral purposes, yet large sections of native mixed-wood forest, wetland, scrub, and riparian habitats still exist, including in protected areas such as Algonquin Provincial Park. We focused on the distribution of four federally listed species-at-risk (SAR) and four species of concern (SOC), consisting of both grassland and forest specialists (Table 1). All have experienced precipitous declines since the 1970s, with population loss in Ontario and Quebec ranging from 55.7% for least flycatcher *Empidonax minimus* to 88.3% for horned lark *Eremophila alpestris* (Table 1; Smith et al. 2019).

**Table 1.** Focal species differentiated by conservation status (SAR – species-at-risk, SOC – species of concern) and habitat specialization (open-country and forest). Population change indicates the average percent decline in Ontario and Quebec from 1970 to 2017 as derived from Breeding Bird Survey data (Smith et al. 2019), while ‘n’ refers to the number of occurrence records for 2018 in the study region.

| Species | Latin name | Status | Habitat | Pop. change | n |
| --- | --- | --- | --- | --- | --- |
| bobolink | *Dolichonyx oryzivorus* | SAR | Open | -84.6% | 421 |
| Eastern meadowlark | *Sturnella magna* | SAR | Open | -86.1% | 323 |
| wood thrush | *Hylocichla mustelina* | SAR | Forest | -72.8% | 195 |
| Eastern wood-pewee | *Contopus virens* | SAR | Forest | -71.0% | 202 |
| horned lark | *Eremophila alpestris* | SOC | Open | -88.3% | 111 |
| Savannah sparrow | *Passerculus sandwichensis* | SOC | Open | -77.0% | 694 |
| killdeer | *Charadrius vociferus* | SOC | Open | -82.0% | 307 |
| least flycatcher | *Empidonax minimus* | SOC | Forest | -55.7% | 158 |

### 2.2. Bird observations

Observations were extracted from the North American Breeding Bird Survey (BBS; Pardieck et al. 2018). BBS surveys have been conducted since 1966 and involve 40 km long transects consisting of 50 road-side point counts separated by ∼0.8 km. Each count lasts 3 min and all birds detected within 400m are recorded. Each route is surveyed once per year during peak breeding season (late-May to early-July) and under fair weather conditions. Rather than summing observations within route segments as is typically done (e.g., Distler et al. 2015), we treated each count as an individual data point to leverage fine-scale habitat associations when predicting species distributions.

We used a single year of BBS data to match remote-sensing land cover estimates collected in 2018 and limit uncertainty introduced by crop rotation. Count data were converted to occurrence data for each species to use in binomial species distribution models. The BBS sampling methodology, combined with one year of data, does not allow estimates of detection probability. To assess the impact of using uncorrected occurrence data, we used an additional validation test based on occurrence data pooled over 5 years (2014–2018; see section 2.5). Due to a large sample size of 5200 points across 104 routes, we expected issues of detectability to have a minimal effect on estimates of species distribution.

### 2.3. Habitat and environmental predictors

For habitat associations, we used 30m resolution land cover data from the 2018 Agriculture and Agri-food Canada annual crop inventory (ISO 19130; Table A1). This data set provides detailed information on up to 71 natural and agricultural land cover classes. For each of a subset of ecologically relevant classes, we calculated the proportional area within a 200m and 1km radius. The resulting 16- and 400-hectare areas allowed us to fit multi-scale species distribution models that incorporated habitat associations at both the territory and landscape scale.

We included the following natural land cover classifications in our analyses: water, wetland, shrub, forest, and grassland. Forests consisted of deciduous (broadleaf), coniferous, and mixed-wood forest cover. For agriculture, we separated land covers into pasture, including grass and forages (e.g., hay), and arable crop (hereafter ‘crop’) following Wilson et al. (2017). Native grassland only represents around 1% of the land cover and therefore it was grouped with pasture. Crops included three main types – corn, soybean, and cereals – as combined they make up >90% of the crop land cover in the region. Finally, we included urban land cover which consists of any human-made structures, such as cities, industrial parks, and roads.

To account for general effects of climate and topography, we extracted breeding season climate variables from the CHELSA Bioclim database and three terrain variables (elevation, slope, aspect) from the Canadian Digital Elevation Model (Table A1). The CHELSA database includes seasonal temperature and precipitation variables averaged from 1901–2016 and thus indicates broad-scale spatial differences in climate. We chose the two most relevant climate variables for the breeding season: mean temperature of the warmest quarter (BIO10), and precipitation of the warmest quarter (BIO18). Across the study region, average breeding season temperature varied by 12.8°C (range: 11.0–23.8°C) and precipitation by 350 mm (170–520mm), with the southwest being warmer and drier on average. From the CDEM, we derived aspect in radians and then converted it from a circular variable using a negative cosine transformation, such that a value of 1 is south, -1 north, and 0 either east or west.

### 2.4. Species distribution models

Habitat suitability was estimated for each species as a function of habitat, terrain, and climate using a 400m resolution grid to match the scale of BBS point counts. For habitat, each grid cell was associated with land cover proportions both within the cell (200m radius), as well as within 1km of the center of the cell. Incorporating multiple spatial scales in species distribution models can outperform single-scale models by accounting for scale-dependent habitat associations and species-specific habitat requirements (Hallman and Robinson 2020). Since point counts were conducted along roads, we included urban cover (which includes road surfaces) at the 1km scale only to avoid overpredicting presence along roadways. For forest-dependent species, we tested models that combined all crop classes into one class ‘crops’ and separated forest cover into its three subclasses: deciduous, coniferous, and mixed-wood. For grassland species, we included forest cover as a general variable and retained individual crop types (cereals, corn, soy) as we expected grassland species to be more responsive to variation in crop type (Table A1).

Prior to model fitting, both presence and absence points were thinned to reduce the potential influence of spatial autocorrelation. We calculated pairwise distances among each point and then randomly removed points until no two were within 1km of each other. This process was replicated 10 times for both presences and absences of each species and the single replicate that retained the most combined data points was selected.

We used ensemble species distribution models (ESDMs) to predict current habitat suitability across the study region for each of the eight focal species. ESDMs integrate multiple modelling approaches, allowing for the modeling of uncertainty among predictions (Thuiller 2004, Elith et al. 2006, Buisson et al. 2010). This is a powerful framework for uncommon species, such as species-at-risk (Breiner et al. 2015), because predictions are informed by multiple techniques that may perform better with certain presence to absence ratios, spatially-structured land cover patterns (i.e., homogenous, heterogeneous), or combinations of habitat predictors (Thuiller 2004, Marmion et al. 2009). We combined Generalized Linear Models (GLMs), General Additive Models (GAMs), and Random Forest (RF) to leverage linear associations, splines, and a decision tree approach, respectively.

Each component model of the ensemble was fit using the same model structure across species. GLMs were fit with all linear and quadratic terms, while GAMS included thin-plate splines with a maximum of 10 knots per variable. RFs were built by growing 5000 trees per replication, with a terminal node size of 1 for fine-scale classification, keeping all other default settings. Each model was replicated 10 times for a total of 30 iterations for each species. All model approaches and replications within the ensemble were integrated using a weighting scheme based on the area under the curve (AUC), such that models with the highest AUC contributed proportionally more to the final prediction. We also set an ensemble threshold of 0.7, where only replicates with an AUC ≥ 0.7 were retained to reduce uncertainty by precluding poorly fit models. The binary threshold which converts habitat suitability to predicted occurrence was estimated by maximizing the True Skills Statistic (TSS; Allouche et al. 2006) while minimizing the difference between sensitivity and specificity (Liu et al. 2005). In this way, we prioritized small omission rates (false absences) rather than commission rates (false presences), which is advisable for uncommon species (Liu et al. 2013). Finally, we estimated variable contribution by calculating the Pearson’s correlation (r_p_) between predictions from the global model and a model with each of the variables sequentially removed.

### 2.5. Model fit and validation

To estimate model fit, we used a hold-out cross validation approach where the data was randomly partitioned into a training and test dataset using a 70:30 split repeated 5 times with replacement for each model replication (i.e., 50 times per model; 150 per species). Model fit was assessed on the test dataset using a combination of AUC, TSS, the omission rate, and the commission rate averaged across replicates. Models with higher AUC or TSS values and low omission/commission rates were considered a better fit. In addition, while BBS data technically has absence data, the survey methodology is not conducive to estimating detectability and thus it is uncertain to what degree locations with undetected species are ‘true’ absences. To validate prediction accuracy and account for imperfect detection, we also calculated commission rate based on occurrence data pooled over 5 years (2014–2018), where a point was only considered a ‘true’ absence if a species went undetected in all years. Using multiple years of data to identify detectability issues has been employed in previous studies using BBS data (e.g., Donovan and Flather 2002, Endenberg et al. 2019). Since we only used one year to fit our models, assessing the accuracy of predicted absences over 5 years allowed us to quantify over-prediction due to issues with model fit or species detectability.

While multi-scale models can be more informative than single-scale models, they run the risk of collinearity and over-fitting. Predictions of species distribution are generally not influenced by collinearity if the collinear structure is consistent in the training and testing data (Dormann et al. 2012, Hallman and Robinson 2020). By evaluating model fit on 150 randomly selected training and testing data sets per species, it is highly unlikely we would consistently sample different collinear structures. However, we tested for over-fitting using two approaches. First, we calculated the difference in AUC and TSS among training and test partitions for each replication (AUC_dif_ and TSS_dif_). High values of AUC_dif_ and TSS_dif_ indicate potential over-fitting and low precision as there is less agreement among partitions. Secondly, we fit fine- and coarse-scale ensemble models to compare with the multi-scale model using the same model fit statistics mentioned above: AUC, TSS, omission rate, and commission rate (1 and 5 years). Either of the reduced models fitting better than the multi-scale model would indicate potential over-fitting (Hallman and Robinson 2020).

### 2.6. Species richness

To evaluate species richness across the study region, we used stacked species distribution models (SSDMs). SSDMs involve overlaying the predicted distribution of each species and summing across layers to estimate species richness, retaining valuable information on species composition (i.e., forest or grassland specialists; Dubuis et al. 2011). While SSDMs can overpredict species richness because species-specific habitat predictors do not reflect macroecological community drivers (e.g., species interactions), the extent of overprediction is often dependent on individual model fit and optimization (Distler et al. 2015, Biber et al. 2019). Therefore, like the focal species, each individual model was optimized to more accurately represent relative species richness across the study region.

We built SSDMS for: 1) grassland specialist SAR and SOC, 2) forest specialist SAR and SOC, 3) all SAR and SOC species combined, and 4) all species that occur within the region (i.e., total species richness). For all species (4), ensemble models were built for all terrestrial birds observed on the BBS routes, excluding non-native species. We also excluded species observed less than 50 times in 2018. Of the 176 species observed in 2018, 76 species met the criteria and were included (Table A2). For consistency with focal species, the same model structure, binary threshold estimation, and fit evaluation were used (sections 2.4–2.5).

Predicted distribution maps were overlaid with those of the 8 focal species and summed to estimate species richness. Variable contribution for species richness was calculated as the average contribution across all species and its associated standard error. We also fit GAMs describing predicted species richness with the land cover, terrain, and climate predictors that were identified as having the greatest variable contribution to evaluate the habitat associations that shape species richness.

### 2.7. Prioritization analysis

We performed a prioritization analysis using Integer Linear Programming (ILP) with the Gurobi optimizer (Gurobi Optimization and LCC, 2020) to evaluate planning scenarios based on our conservation objective. Our goal was to maximize the effectiveness of limited conservation funds by optimizing the trade-off between the conservation of all targets and socioeconomic cost. In this case, the targets refer to the distributions of the 8 focal species and the upper 25% of areas with the highest total species richness. We used a minimum set objective with the goal of retaining at least 25% representation of all targets in the most cost-effective manner by minimizing the number of planning units needed to reach this goal. The planning units for the prioritization were 2 km^2^ terrestrial pixels (1km radius). Distribution maps for each target were created at a 0.4 km^2^ scale (200m radius), but each one was individually aggregated to a 2 km^2^ scale for the prioritization as the more likely scale at which lands are managed. Portions of the study area that were already formally protected were locked into the solution. Thus, we identified additional non-protected areas that, when combined with existing protected areas, met the minimum set objective.

A conceptual diagram of all modelling steps is including in Figure 1B. All spatial and statistical analyses were conducted in R software version 3.6.3 (R Core team 2020). GLMs were fit using package ‘lme4’ (Bates et al. 2015), GAMS with ‘mgcv’ (Wood 2011), and RFs with ‘randomForest’ (Liaw and Wiener 2002). Individual species distribution models were combined into ESDMs using a modified version of ‘SSDM’ (Schmitt et al. 2017). The prioritization analysis was conducted using ‘prioritizr’ (Hanson et al. 2020).

## 3. Results

### 3.1. Predicted distributions of focal species

In 2018, at least one of the eight focal species was observed at approximately 30% of the 5200 BBS point counts. The number of observations per species ranged from a low of 111 for horned lark to a high of 694 for Savannah sparrow (Table 1). Model fit was generally high, with an average AUC of 0.860 (range: 0.826–0.921) and TSS of 0.732 (range: 0.653–0.869). The rate of false negatives (omission error) was also low and within expected ranges (0.030–0.184) as was the false positive or commission error rate (0.101–0.199), indicating high predictive accuracy across species (Table A3). Model comparisons among the multi-, fine-, and coarse-scale models clearly indicated an advantage to using a multi-scale approach as it was the best fitting model across all species (Table A3). Predictions within each ensemble model were moderately to highly correlated (r_p_ > 0.70), suggesting each component model was capturing similar relationships, except for the three forest specialists where RF and GLM were in poor agreement, highlighting the value of an ensemble approach (r_p_ ∼ 0.50; Table A4).

As expected, predicted distributions indicated grassland specialists have more suitable habitat in the southern and southeastern regions of Ontario where land cover is dominated by agriculture, although substantial areas of suitability were also predicted along the St. Lawrence River in Quebec (Figure 2). Forest specialists concentrated in the northern portions of the study region, yet suitable habitat was still identified within the more intensive agricultural area in the south, particularly near Lake Huron which represents a mosaic of field crops, forest patches, and hedgerows (Figure 2). Grassland (including pasture, forage, and hay) had the strongest influence on the distribution of bobolink, Eastern meadowlark, and Savannah sparrow, but had little influence on horned lark and killdeer (Table A5). Instead, these latter two species responded positively to row crops, particularly cereals but also soybean for horned lark (Table A5). Forest specialists primarily responded to forest availability, with particularly strong and positive associations with deciduous, and for wood thrush, mixed-wood forest cover (Table A5). The influence of scale also differed among species, highlighting the importance of a multi-scale approach. For example, while grassland species were generally associated with grassland and agriculture land-use at the territory scale (200m radius), horned lark responded most strongly to agricultural land cover at larger spatial scales (1km; Table A5).

**Figure 2.**
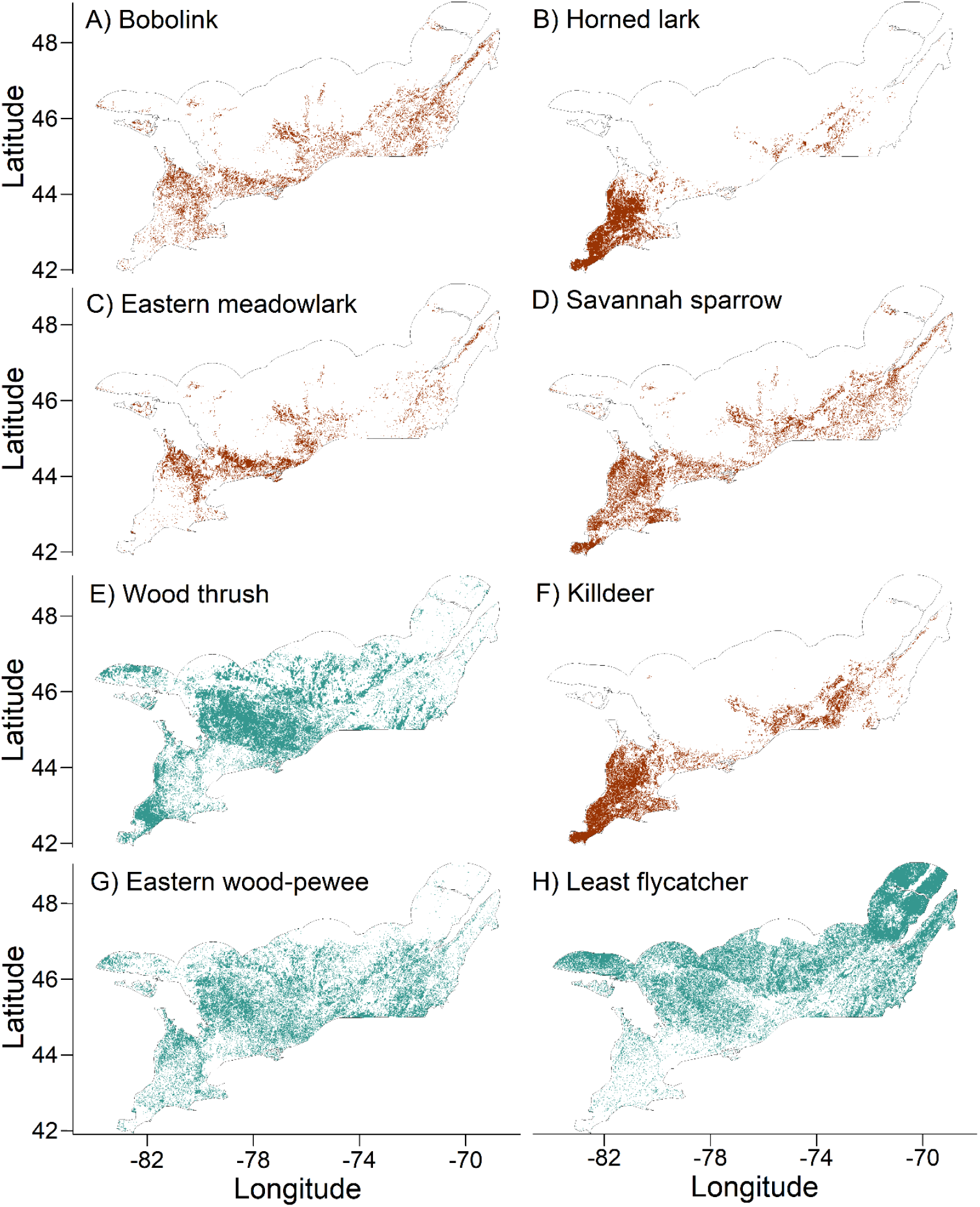
Estimates of species distribution or the fundamental niche of the four species-at-risk (a, c, e, g) and species of concern (b, d, f, h) based on the estimated habitat suitability threshold. Distributions in brown and green indicate grassland and forest specialists, respectively.

### 3.2. Focal species richness

For the focal at-risk community, our models predicted higher grassland specialist richness in the south and southwest, whereas greater forest specialist richness occurred in the north and northwest (Figure 3), paralleling results from individual focal species models. The proportion of grassland within a 200m radius had the greatest influence on grassland species richness, followed by moderately strong influences of row crops at both 200m and 1km radii (Figure 3), again, aligning with results from our focal species. Forest species richness was driven primarily by deciduous and mixed-wood forest cover, as well as more moderate influences of wetland and shrub at larger spatial scales (Figure 3

**Figure 3.**
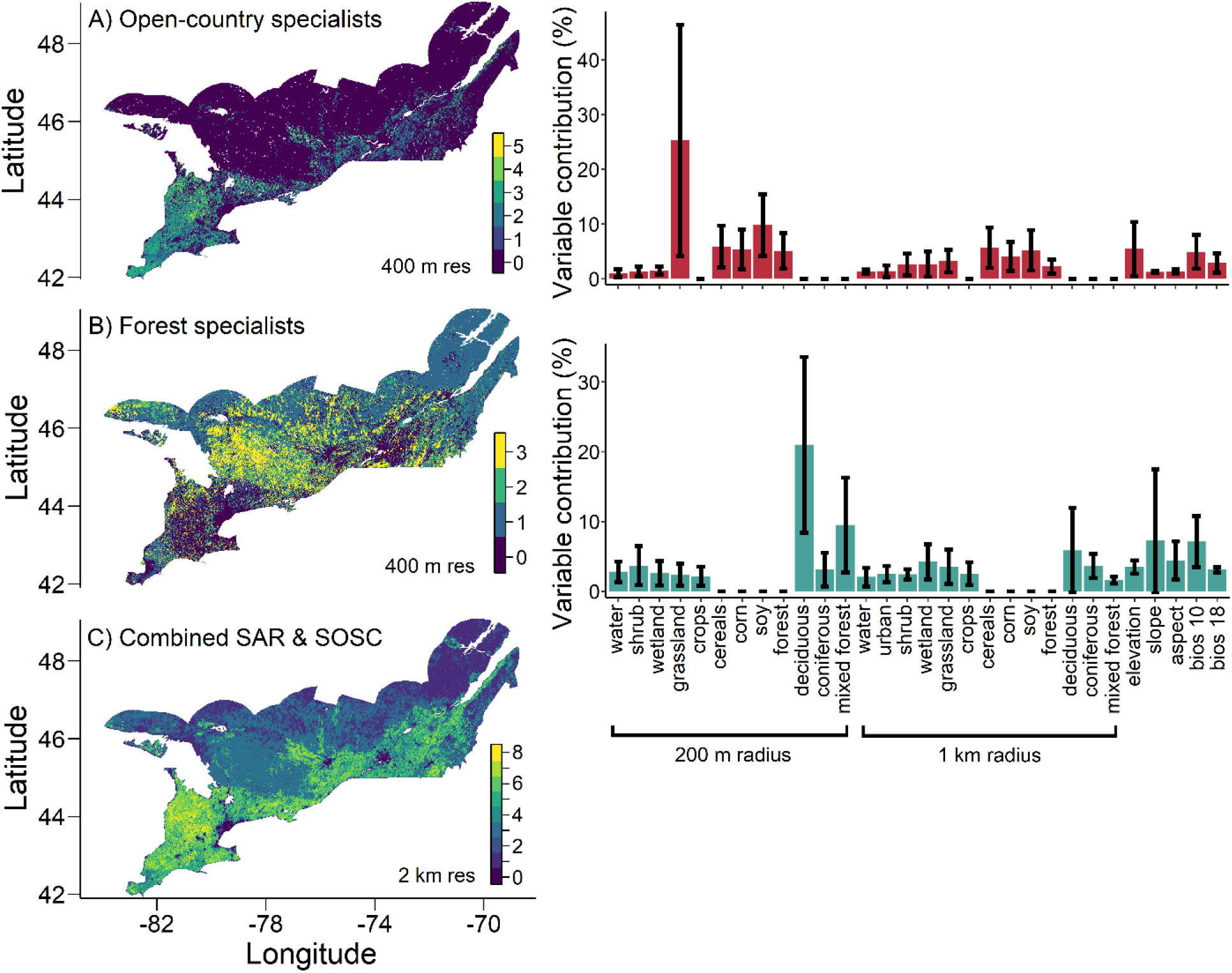
Predicted richness of focal species richness, depicting: A) grassland specialists only, B) forest specialists only, and C) all focal species combined. For (C), the predicted distributions of open-country and forest specialists were summed within 2 km^2^ (1km radius) to demonstrate the richness of focal species at the landscape scale rather than the territory scale. The adjacent bar plots indicate average variable contribution as a percentage for each of the grassland and forest specialists. Error bars depict standard deviation among component species.

Combined richness of at-risk grassland and forest communities was greatest to the west and northwest of Toronto, near Lake Huron, as well as west, southwest of Montreal and Ottawa (Figure 3C). Again, these are regions with greater habitat heterogeneity, including fields of pasture or crop interspersed with forest and woody shrub patches. This was reflected in the top habitat associations, where at-risk species richness increased with the proportion of shrub at the territory scale and wetland at the broader landscape scale (Figure 4). Importantly, the positive influence of grassland and deciduous forest both peaked around 40–50% of land cover, indicating that a mix of habitat types and forest edges maximizes diversity of all focal species (Figure 4). At the broader scale, the influence of grassland and corn also plateaued around 40% of land cover, while cereals had a strong linear effect, likely driven by the positive influence on species like horned lark, Savannah sparrow, and killdeer (Figure 4). Temperature (BIO10) and precipitation (BIO18) had positive and negative influences on richness, respectively, suggesting warmer and moderately dry climates are favoured, although the magnitude of influence was significantly less than the landcover effects (Figure 4).

**Figure 4.**
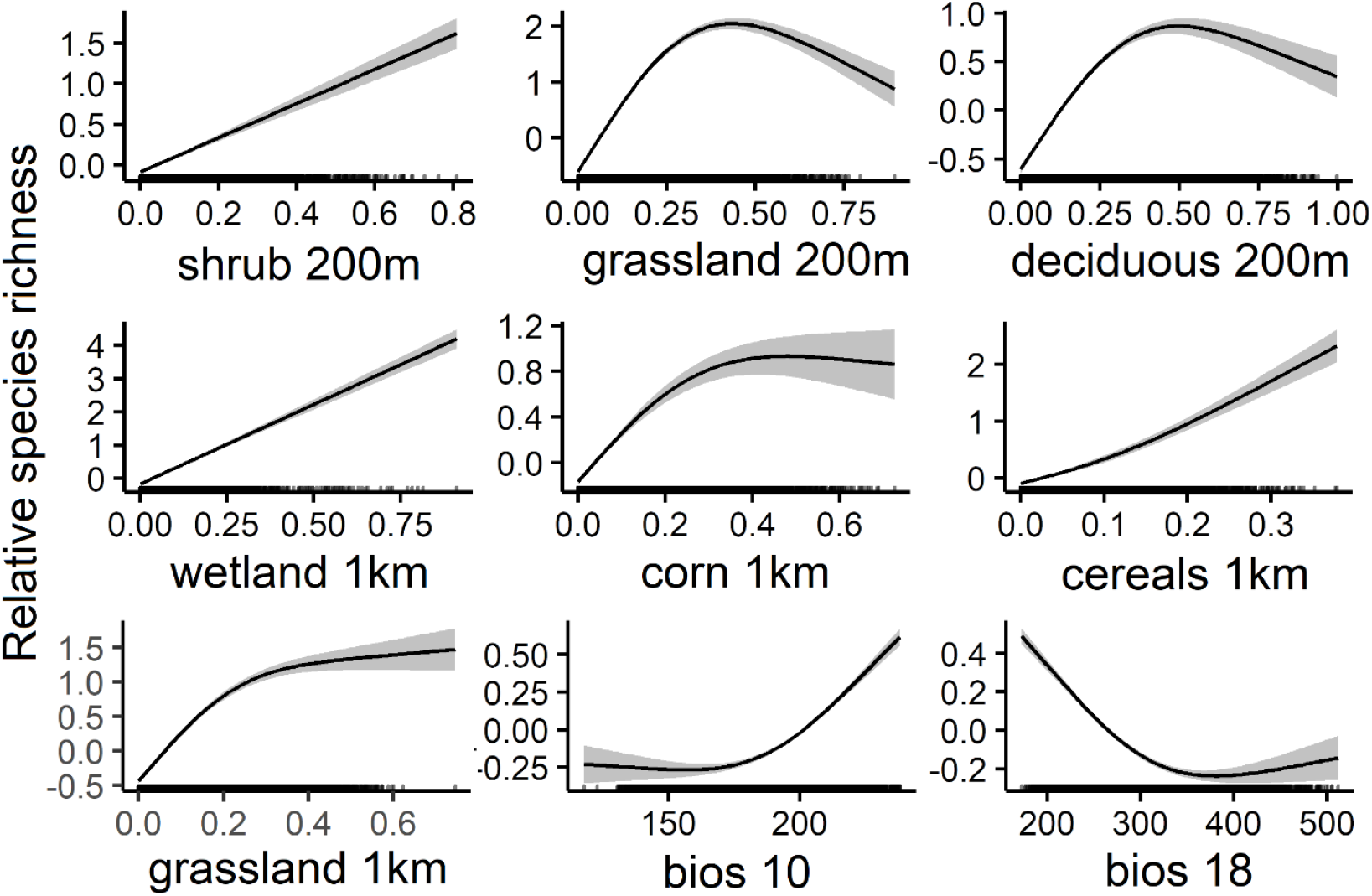
The top habitat associations predicting focal species richness. Relative species richness is the difference from the mean species richness at zero. For all associations, the X-axis is proportion of land cover, except for bios 10 (temperature) and bios 18 (precipitation) which are displayed in 1/10^th^ degrees Celsius and mm, respectively. The shaded band represents the 95% CI and the distribution of points is depicted along the X-axis.

### 3.3. Overall species richness

Species richness of the overall bird community (76 species) was predicted to reach ∼50–60 species in multiple mid-latitude areas from east to west (Figure 5A). We quantified areas with high species richness, defined as containing at least 75% of the total species richness or 75% of the focal species. This demonstrated that greater than average richness for both groups was predicted in regions near Lake Huron, west of Montreal and Ottawa, as well as along the St.Lawrence River (Figure 5B). These regions contain the necessary land cover, terrain, and climate features that jointly facilitate high species richness of declining species, as well as the overall grassland and forest bird communities.

**Figure 5.**
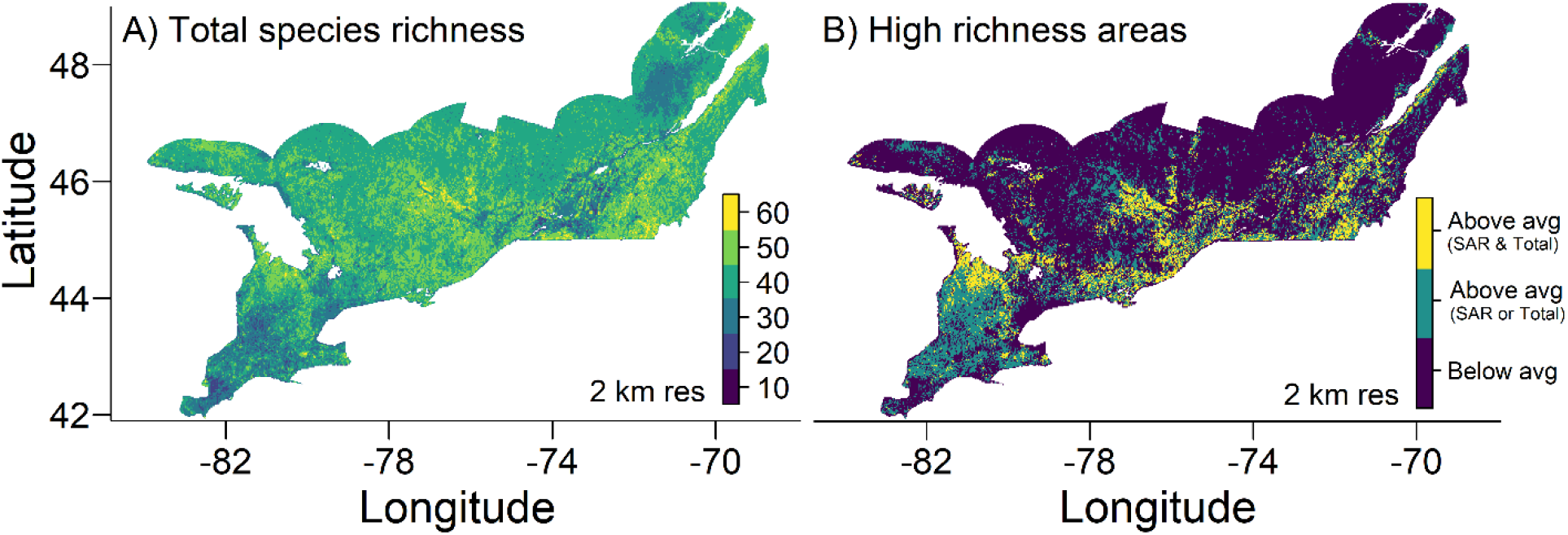
Total species richness predicted within the study region, showing: A) total richness of all terrestrial birds with at least 50 observations (76 species), and B) areas with richness greater than the 3^rd^ quantile (>75% of maximum). For the latter, green indicates areas where species richness is greater than the 3^rd^ quantile for either the focal species-at-risk (≥ 6 species) or total species richness (≥ 41 species), while yellow depicts areas where high species-at-risk and total richness align.

### 3.4. Prioritization analysis

The minimum set objective required 16.7% (64,800 km^2^) of the focal land area of which 48% (31,040 km^2^) lacked existing protection (Figure 6). The prioritized and protected areas were largely concentrated in the northern portions of the focal area with considerable blocks of protected area represented by larger parks (e.g., Algonquin Provincial Park in Ontario, Mont-Tremblant National Park in Quebec). The prioritized and unprotected area included many small parcels, mostly in the south with larger contiguous selections in the southwest, near Lake Huron, and south of the St. Lawrence River Valley in the east (Figure 6). While the minimum set objective ensured 25% of each species’ distribution was included, variable distribution extents among species translated into considerable differences in selected areas ranging from 6,068 km^2^ for Eastern meadowlark to 42,992 km^2^ for least flycatcher (Table A6).

**Figure 6.**
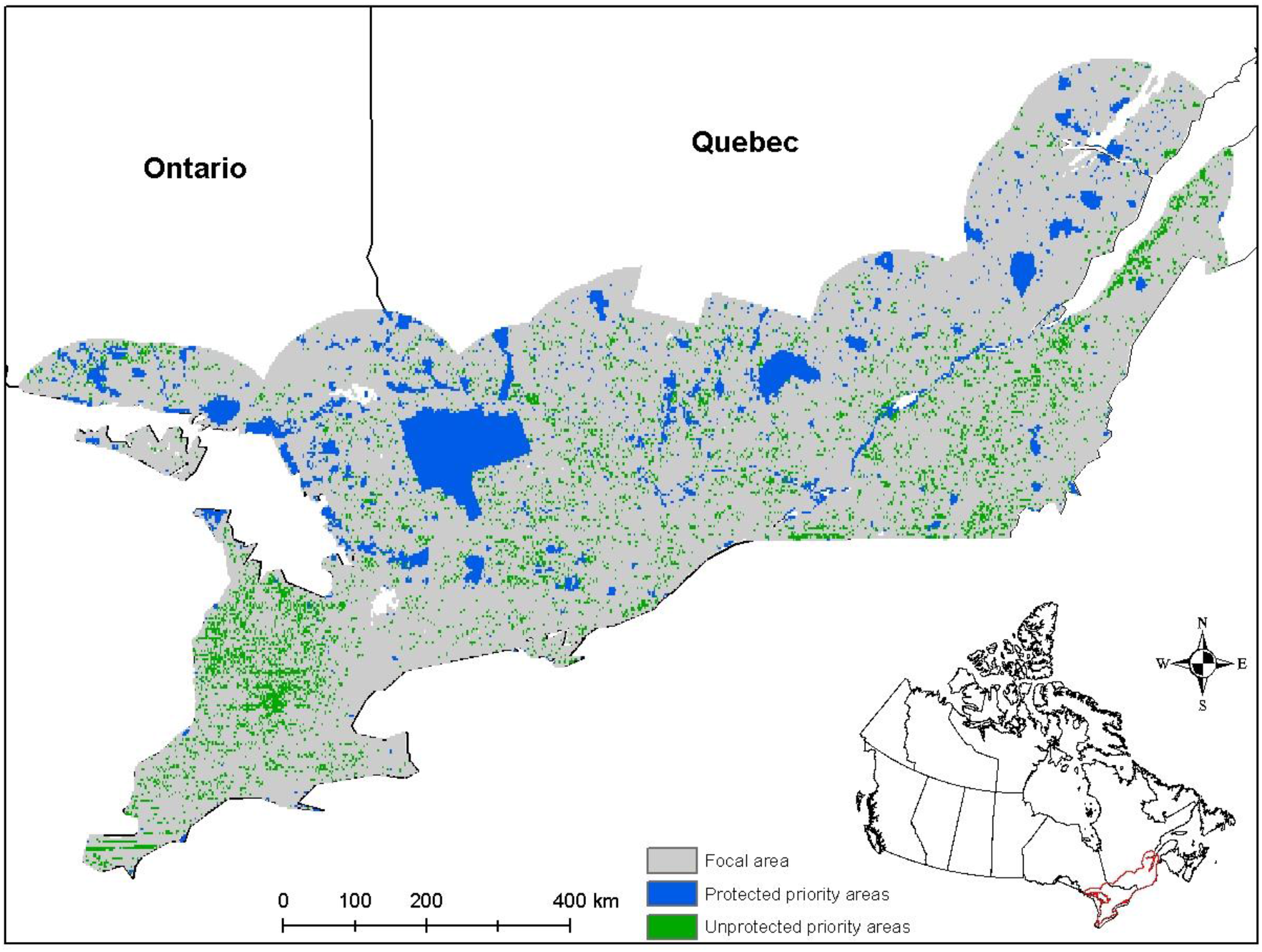
Minimum set prioritization results showing selected sites with existing protection (blue) and without protection (green). The prioritization selected at least 25% of the distribution for each feature in the most cost-effective manner (i.e., minimum required area). Features were the distribution maps for each species shown in Figure 3 and the upper quartile for total species richness shown in Figure 5A.

## 4. Discussion

Conservation planning frequently requires trade-offs to meet multiple objectives. In this study, we examined how to balance the conservation needs of grassland and forest birds within a naturally forested ecosystem where the expansion of agriculture has resulted in a vastly different ecosystem with consequent shifts in bird communities (Wilson et al. 2020). Despite extensive agriculture land-use, we were still able to identify areas and landscape types that were suitable for individual species-at-risk and maximized the diversity of bird communities dependent on grassland and forest. Habitat associations and the spatial scale at which they influenced occupancy differed greatly among individual focal species and indicated the importance of mixed-wood forests, wetland, native grassland and pastoral agriculture (i.e., forage, hay), as well as certain row crops such as cereals which enhanced occupancy of some grassland species of concern. Despite differences in habitat requirements among species, several regions characterized by complex landscape structures were predicted to balance the needs of both grassland and forest-associated species.

Our study highlights the importance of maintaining habitat heterogeneity in agroecosystems, a result that has been shown across different temperate and tropical ecosystems globally (Karp et al. 2012, Liu et al. 2013). Areas with high species richness and selected by the prioritization included two types of heterogeneity. In western portions of the study region south of Lake Huron, the landscape is largely agricultural but with considerable representation of linear woody features along roadways and between fields (Figure S1). These features include hedgerows and riparian strips associated with natural and drainage ditch networks, which have been shown to enhance the diversity of forest and shrub-bird communities in both eastern Canada (Wilson et al. 2017) and elsewhere (Hinsley and Bellamy 2000, Sykes and Hannon 2001, Bátary et al. 2010, Mulwa et al. 2012). Richness and the number of prioritized sites declined to the south of this region near Lake Erie where minimal woody cover has been retained (Figure S1). Parts of the eastern study region (e.g., south and east of Montreal) also included areas of high richness that were selected by the prioritization. In contrast to the areas south of Lake Huron, where habitat heterogeneity was maintained via linear strips within a largely agricultural region, the areas south of the St. Lawrence Valley in Quebec were often a combination of forest blocks and open agricultural fields, thus highlighting two different types of landscapes that can meet the conservation challenge we addressed in this study.

As forests are the primary land cover in the region, the potential distribution of forest specialists tended to be considerably greater than that for grassland species. When using a minimum set prioritization with a goal to protect a percentage of each species distribution, the result is a higher area target for forest species. For example, a 25% area goal required only 6,068 km^2^ for Eastern meadowlark, but 42,992 km^2^ for least flycatcher. Because much of the northern portion of the study region contains little agriculture and most of the forested protected areas, this higher area goal for forest species could be met more easily by utilizing the existing protected area network and incorporating forested provincial crown land in management strategies. In contrast, the more challenging goal will be to meet area targets in high value agricultural regions where almost all land is private and strictly protected areas are often not feasible; yet these are the regions needed to reach the area targets for grassland species while also accommodating forest species. In these cases, the objective should be to work with landowners to manage landscapes in a manner that maximizes the diversity of both avian groups to the extent possible without compromising agricultural yields.

It is important to note that we assessed fundamental habitat suitability, or the potential distribution that each focal species could inhabit, rather than the realized distribution (Hutchinson 1957, Pulliam 2000). Various factors can limit realized relative to fundamental distribution, including 1) avoidance of certain sites due to human activity (e.g., recreation), and 2) species interactions, which were not addressed in this study. Additionally, 3) legacy effects may be particularly important with agriculture as crop rotation can change land cover from year to-year and thus settlement decisions of philopatric individuals may be partially influenced by previous land cover types. Finally, 4) population size may be limited by non-breeding season effects stemming from different periods across the annual cycle (Taylor and Stutchbury 2016, Wilson et al. 2018), such that suitable habitat may remain unoccupied. We found relatively low commission error rates (false positives) even when pooling across 5 years, suggesting our models were accurately differentiating between good and poor breeding habitat. That said, wood thrush, least flycatcher, and eastern wood-pewee had the highest 5-year commission rates (14–18%). Since these are Neotropical migrants where non-breeding season threats are now well established, it is possible that these higher commission rates are partially explained by poor survival during migration and on the wintering grounds, leading to lower occupancy rates of otherwise suitable habitat.

Future habitat suitability predictions can be further improved by integrating different habitat features. We focused primarily on the proportion of various land covers to describe habitat suitability for each species. Our results highlighted the importance of diverse habitat complements across multiple scales but did not shed any light on the relative position of certain habitat types within either territory or landscape scales which may impact habitat suitability (e.g., patch size, structural connectivity; Andrén 1994, Mortelliti et al. 2010, Fahrig 2020). Future research should refine predictions by incorporating variables such as habitat aggregation, patch size, and structural connectivity (i.e., placement of linear woody features) to improve recommendations on how to manage for a more diverse agroecosystem.

### 4.1. Application of bird conservation in an agroecosystem

Our results provide a framework to derive habitat management priorities focused on preserving both forest and grassland bird communities in southern Ontario and Quebec, a region that is representative of many intensive agroecosystems in north temperate regions. We expect this information could be used in programs such as Canada’s Habitat Stewardship Program, which allows agencies to identify areas and enact stewardship programs for species-at-risk recovery. Retaining strips of woody habitat as a strategy to maintain biodiversity may be particularly suitable in landscapes with higher agriculture value, requiring minimal agricultural land while providing numerous ecosystem services such as limiting soil erosion, windbreaks to minimize crop damage, riparian corridors for reducing runoff pollution, and supporting insect and faunal communities that provide pest control and pollination (Marshall and Moonen 2002, Swinton et al. 2007, Tscharntke et al. 2012). Retaining forest blocks may be more feasible in regions with lower agricultural value, occurring more often when grassland/pasture represented a high proportion of agricultural land-use in our study region. With little remaining grassland, these landscapes allow grassland-dependent species to persist, including federally listed species such as Eastern meadowlark and bobolink (Wilson et al. 2017). Therefore, in addition to crop type, actions benefitting grassland-dependent species also need to target favourable agricultural practices, including reduced fertilizer and pesticide inputs, multi-year forage production within corn-soy rotations to provide consistency in preferred forage cover types, and limiting mortality of offspring by scheduling hay harvest to reduce overlap with peak breeding season (Put et al. 2020).

## Authors’ Contributions

All authors contributed to study design. DDZ conducted the analyses with assistance from NA on land cover assembly and extraction. DDZ and SW co-led writing of the manuscript with feedback and revision from all authors.

## Data Accessibility

Data and code will be made available in a public repository upon manuscript acceptance.

Conflict of Interest

The authors declare no conflict of interest.

## Acknowledgements

We thank all volunteers who collect the Breeding Bird Survey data that allowed us to conduct this analysis. Funding for this work was provided by the Interdepartmental Initiatives in Agriculture Grant of Agriculture and Agri-Food Canada.

## Appendix

**Table A1.** Overview of habitat predictors for species distribution models (SDMs). Guild indicates the species group associated with the predictor (G = grassland, F = forest). ‘Original res’ is the resolution at which the raster layer was available, while ‘Res’ is the resolution incorporated into the model and ‘Method’ indicates the aggregation/resampling method by which the resolution was converted.

| Variable | Guild | Original res | Res | Method | Description | Source |
| --- | --- | --- | --- | --- | --- | --- |
| *Natural land cover* |  |  |  |  |  |  |
| Water | G/F | 30m | 400m<br>2km | Sum | Lakes, rivers, streams | AAFC<sup>1</sup> |
| Shrub | G/F | 30m | 400m<br>2km | Sum | Low-profile woody vegetation (+/- 2m height) | AAFC<sup>1</sup> |
| Wetland | G/F | 30m | 400m<br>2km | Sum | High water table with semi-permanent/permanent wetland veg (eg, marsh, bog, swamp) | AAFC<sup>1</sup> |
| Forest | G | 30m | 400m<br>2km | Sum | Coniferous, deciduous, & mixed-wood forest. | AAFC<sup>1</sup> |
| Deciduous | F | 30m | 400m<br>2km | Sum | Broadleaf forest only. | AAFC<sup>1</sup> |
| Coniferous | F | 30m | 400m<br>2km | Sum | Coniferous forest only. | AAFC<sup>1</sup> |
| Mixed-wood | F | 30m | 400m<br>2km | Sum | Mixed deciduous & coniferous forest or treed areas. | AAFC<sup>1</sup> |
| Grassland | G/F | 30m | 400m<br>2km | Sum | Native grassland (grasses, herbaceous veg), pasture, & forage (periodically cultivated grasses and perennial crops such as clover grown for hay, pasture, or seed). | AAFC<sup>1</sup> |
| *Anthropogenic land cover* |  |  |  |  |  |  |
| Crops | F | 30m | 400m<br>2km | Sum | Combination of cereals, corn, and soybean (>90% of agricultural land). | AAFC<sup>1</sup> |
| Cereals | G | 30m | 400m<br>2km | Sum | Consists of barley, millet, oats, rye, spelt, triticale, switchgrass, sorghum, quinoa, and winter/spring wheat. | AAFC<sup>1</sup> |
| Corn | G | 30m | 400m<br>2km | Sum | Predominantly corn crops. | AAFC<sup>1</sup> |
| Soybean | G | 30m | 400m<br>2km | Sum | Predominantly soybean crops. | AAFC<sup>1</sup> |
| Urban | G/F | 30m | 2km | Sum | Developed land with associated veg (eg, roads, cities, industrial sites, golf courses, etc.) | AAFC<sup>1</sup> |
| *Topography* |  |  |  |  |  |  |
| Elevation | G/F | 22m | 400m | Avg | Height above sea level (m). | CDEM<sup>2</sup> |
| Slope | G/F | 22m | 400m | Avg | Incline (°) calculated from adjacent elevation. | CDEM<sup>2</sup> |
| Aspect | G/F | 22m | 400m | Avg | North-south orientation | CDEM<sup>2</sup> |
| *Climate* |  |  |  |  |  |  |
| Bios10 | G/F | 1km | 400m | Bilinear | Mean temp of warmest quarter | CHELSEA<sup>3</sup> |
| Bios18 | G/F | 1km | 400m | Bilinear | Precipitation of warmest quarter | CHELSEA<sup>3</sup> |

<sup>1</sup>Annual crop inventory assembled by the Earth Observation Team of the Science and Technology Branch of Agriculture and Agri-food Canada (AAFC; Government of Canada; Access: https://open.canada.ca/data/en/dataset/ba2645d5-4458-414d-b196-6303ac06c1c9).
<sup>2</sup>Canadian Digital Elevation Model, 1945-2011, Canada Center for Mapping and Earth Observation, Natural Resources Canada, Government of Canada (Accessed: https://open.canada.ca/data/en/dataset/7f245e4d-76c2-4caa-951a-45d1d2051333).
<sup>3</sup>CHELSEA Bioclim data (Karger et al. 2017a,b)

**Table A2.**
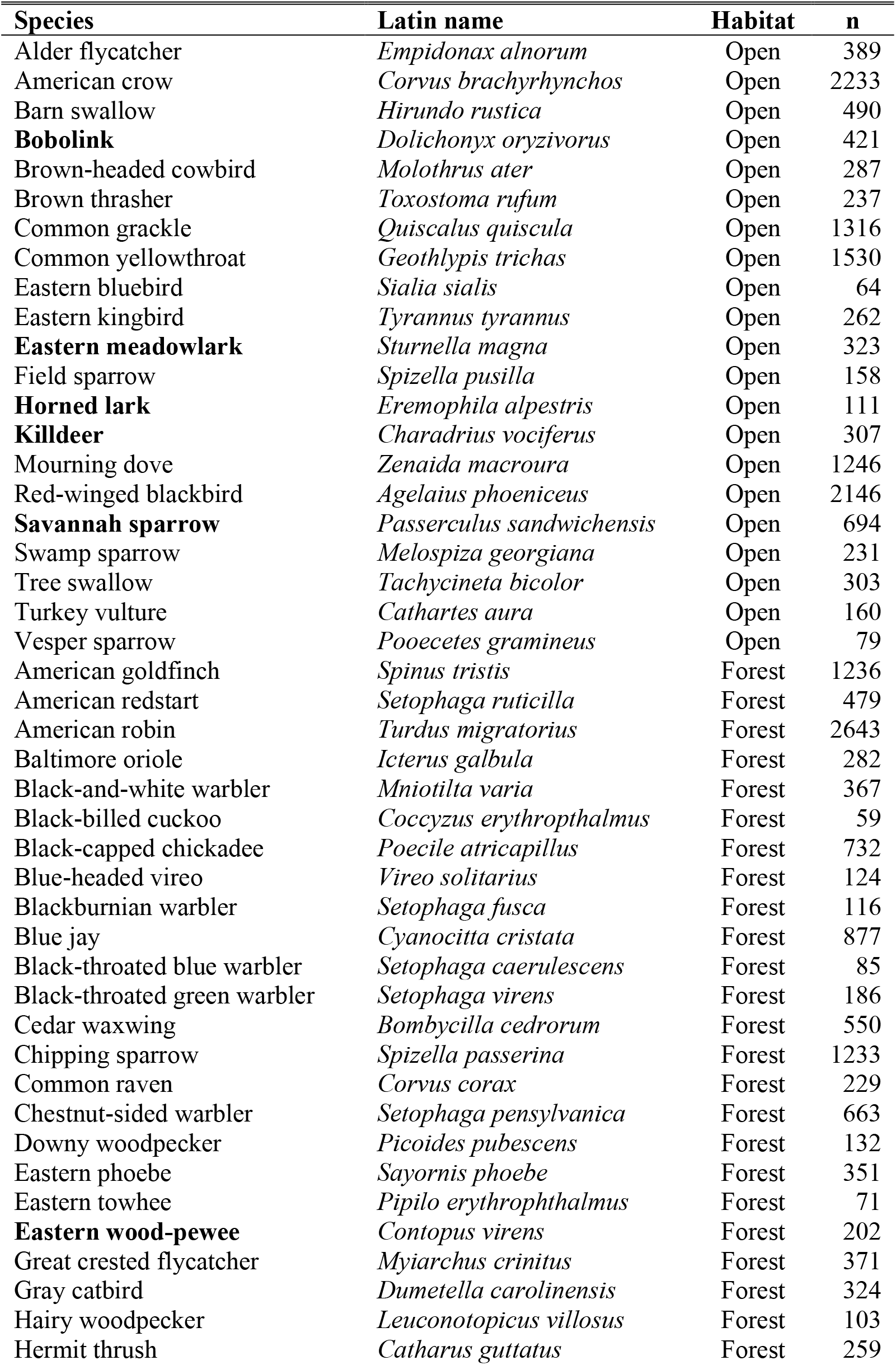
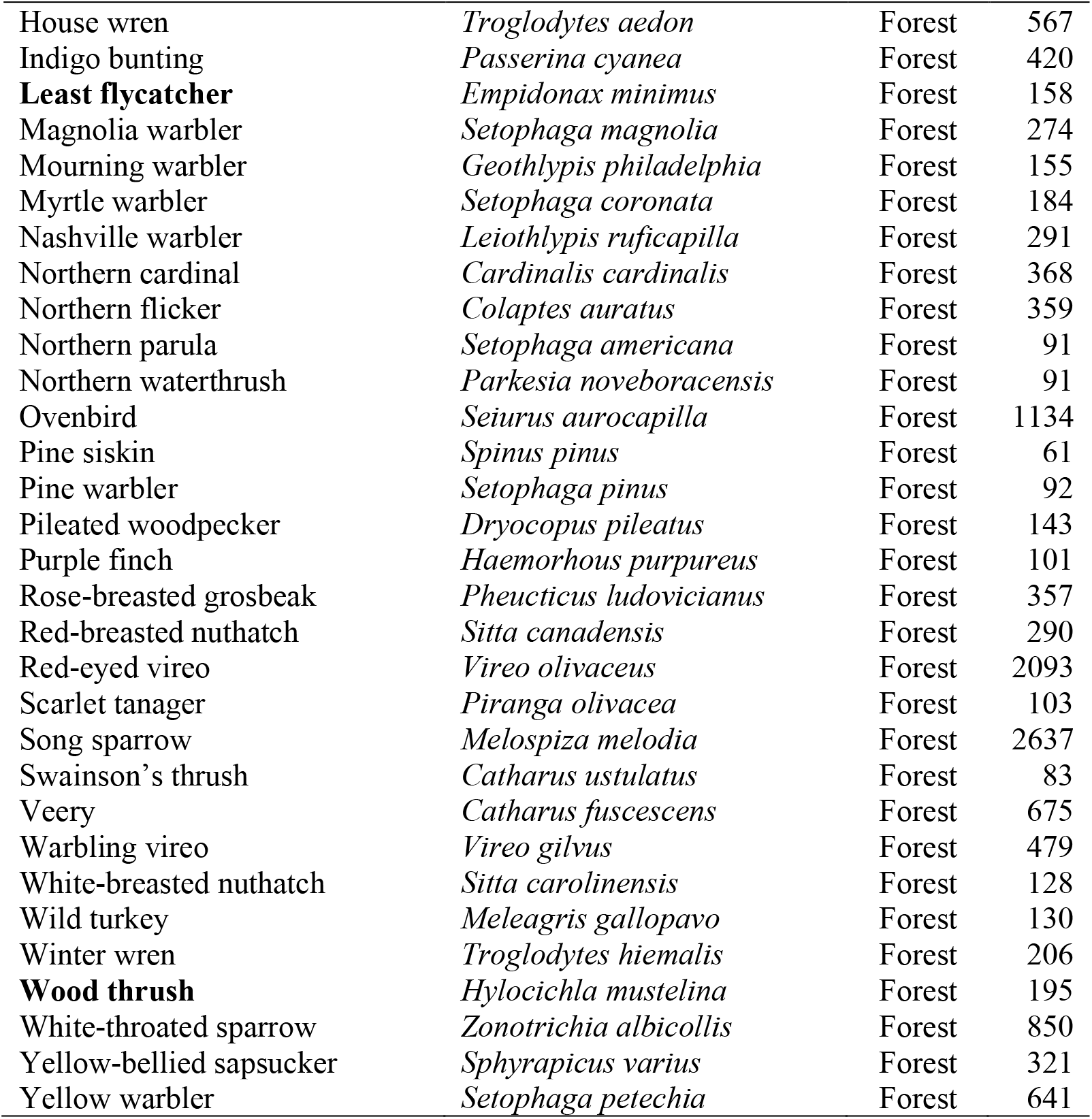
Species included in the stacked species distribution models (SSDMs) for assessing species richness. Observations (n) are the number of sites each species was observed at in 2018. Open refers to grassland-dependent species. All focal species are highlighted in bold.

**Table A3.** Model fit metrics for each focal species comparing the multi-scale model to the fine and coarse-scale models. The model highlighted in bold was considered the top model based on the AUC, TSS, omission rate (OR; false absences) and commission rate (CR; false presences). The commission rate was also calculated over 5 years (2014–2018) to account for imperfect detection. Accuracy is the proportion of correctly predicted presences and absences combined. AUC_dif_ and TSS_dif_ are the average differences between training and testing AUC and TSS, respectively.

| Model | AUC | TSS | OR | CR | CR (5yr) | Accuracy | $AUC_{dif}$ | $TSS_{dif}$ |
| --- | --- | --- | --- | --- | --- | --- | --- | --- |
| *Bobolink* |  |  |  |  |  |  |  |  |
| **Multi** | 0.852 | 0.711 | 0.089 | 0.199 | 0.138 | 0.815 | 0.01 | 0.02 |
| Fine | 0.847 | 0.701 | 0.068 | 0.230 | 0.172 | 0.791 | 0.04 | 0.04 |
| Coarse | 0.820 | 0.646 | 0.098 | 0.255 | 0.198 | 0.765 | 0.02 | 0.01 |
| *Eastern meadowlark* |  |  |  |  |  |  |  |  |
| **Multi** | 0.881 | 0.774 | 0.060 | 0.166 | 0.106 | 0.844 | 0.02 | 0.03 |
| Fine | 0.849 | 0.709 | 0.088 | 0.203 | 0.145 | 0.808 | 0.01 | 0.05 |
| Coarse | 0.836 | 0.681 | 0.092 | 0.227 | 0.155 | 0.787 | 0.04 | 0.08 |
| *Horned lark* |  |  |  |  |  |  |  |  |
| **Multi** | 0.921 | 0.869 | 0.030 | 0.101 | 0.052 | 0.902 | 0.02 | 0.09 |
| Fine | 0.912 | 0.850 | 0.040 | 0.110 | 0.072 | 0.893 | 0.01 | 0.23 |
| Coarse | 0.888 | 0.801 | 0.090 | 0.109 | 0.066 | 0.892 | 0.02 | 0.21 |
| *Savannah sparrow* |  |  |  |  |  |  |  |  |
| **Multi** | 0.839 | 0.691 | 0.122 | 0.187 | 0.119 | 0.826 | 0.01 | 0.02 |
| Fine | 0.835 | 0.682 | 0.120 | 0.198 | 0.122 | 0.817 | 0.01 | 0.11 |
| Coarse | 0.809 | 0.629 | 0.122 | 0.249 | 0.173 | 0.776 | 0.02 | 0.14 |
| *Killdeer* |  |  |  |  |  |  |  |  |
| **Multi** | 0.870 | 0.758 | 0.083 | 0.159 | 0.100 | 0.848 | 0.01 | 0.05 |
| Fine | 0.835 | 0.687 | 0.067 | 0.246 | 0.165 | 0.772 | 0.01 | 0.05 |
| Coarse | 0.807 | 0.630 | 0.103 | 0.267 | 0.183 | 0.750 | 0.03 | 0.03 |
| *Wood thrush* |  |  |  |  |  |  |  |  |
| **Multi** | 0.866 | 0.735 | 0.118 | 0.148 | 0.139 | 0.855 | 0.05 | 0.04 |
| Fine | 0.821 | 0.641 | 0.094 | 0.264 | 0.265 | 0.747 | 0.05 | 0.15 |
| Coarse | 0.803 | 0.606 | 0.141 | 0.253 | 0.228 | 0.755 | 0.04 | 0.18 |
| *Eastern wood-pewee* |  |  |  |  |  |  |  |  |
| **Multi** | 0.826 | 0.662 | 0.152 | 0.186 | 0.174 | 0.817 | 0.01 | 0.03 |
| Fine | 0.771 | 0.550 | 0.103 | 0.346 | 0.338 | 0.672 | 0.04 | 0.10 |
| Coarse | 0.776 | 0.561 | 0.141 | 0.298 | 0.305 | 0.714 | 0.03 | 0.16 |
| *Least flycatcher* |  |  |  |  |  |  |  |  |
| **Multi** | 0.828 | 0.653 | 0.184 | 0.163 | 0.177 | 0.836 | 0.03 | 0.07 |
| Fine | 0.768 | 0.530 | 0.136 | 0.334 | 0.356 | 0.678 | 0.01 | 0.20 |
| Coarse | 0.779 | 0.552 | 0.143 | 0.305 | 0.330 | 0.704 | 0.01 | 0.20 |

**Footnote:** Small differences in AUC and TSS among data partitions ( $AUC_{dif} = 0.01 - 0.05$ ; $TSS_{dif} = 0.02 - 0.09$ ) suggest high consistency among partitions and no indication of over-fitting. Consistent selection of the multi-scale model as the best fit model also suggests that any collinearity among scales is not leading to systematic issues with over-fitting.

**Table A4.** Correlation among predictions of each component model within the ensemble species distribution models for the focal species.

|  | Model | Correlation |  |  |
| --- | --- | --- | --- | --- |
|  |  | GLM | GAM | RF |
| *Bobolink* |  |  |  |  |
|  | GLM | 1.00 | — | — |
|  | GAM | 0.94 | 1.00 | — |
|  | RF | 0.81 | 0.82 | 1.00 |
| *Eastern meadowlark* |  |  |  |  |
|  | GLM | 1.00 | — | — |
|  | GAM | 0.87 | 1.00 | — |
|  | RF | 0.81 | 0.80 | 1.00 |
| *Horned lark* |  |  |  |  |
|  | GLM | 1.00 | — | — |
|  | GAM | 0.80 | 1.00 | — |
|  | RF | 0.82 | 0.82 | 1.00 |
| *Savannah sparrow* |  |  |  |  |
|  | GLM | 1.00 | — | — |
|  | GAM | 0.88 | 1.00 | — |
|  | RF | 0.82 | 0.84 | 1.00 |
| *Killdeer* |  |  |  |  |
|  | GLM | 1.00 | — | — |
|  | GAM | 0.83 | 1.00 | — |
|  | RF | 0.81 | 0.73 | 1.00 |
| *Wood thrush* |  |  |  |  |
|  | GLM | 1.00 | — | — |
|  | GAM | 0.65 | 1.00 | — |
|  | RF | 0.57 | 0.71 | 1.00 |
| *Eastern wood-pewee* |  |  |  |  |
|  | GLM | 1.00 | — | — |
|  | GAM | 0.91 | 1.00 | — |
|  | RF | 0.55 | 0.52 | 1.00 |
| *Least flycatcher* |  |  |  |  |
|  | GLM | 1.00 | — | — |
|  | GAM | 0.76 | 1.00 | — |
|  | RF | 0.49 | 0.60 | 1.00 |

**Table A5.**
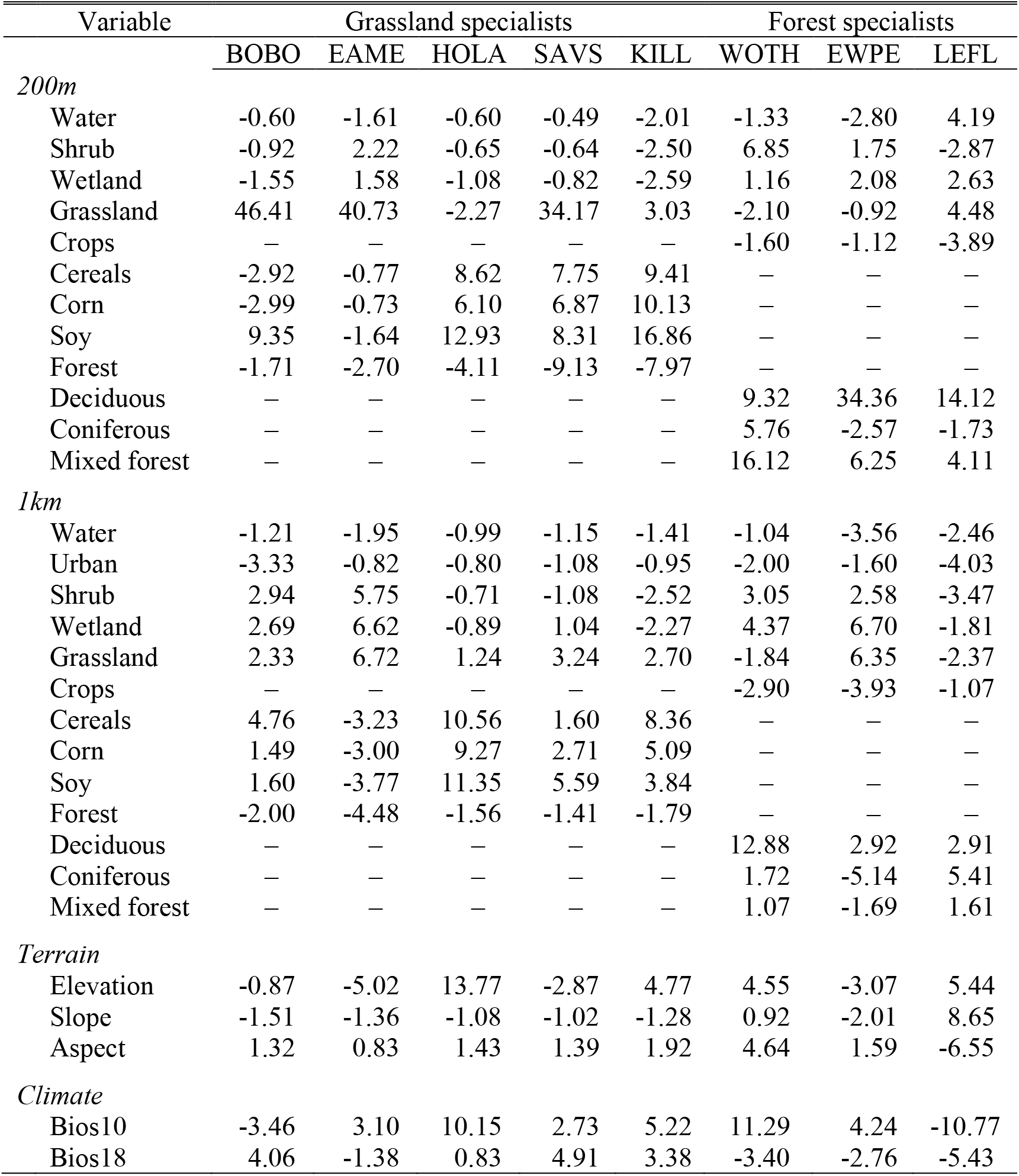
Variable importance for each focal species as derived from the ensemble species distribution models. A dash means that the association is negative, while no sign indicates a positive association. Bios 10 and Bios 18 refer to temperature and precipitation during the breeding season, respectively.

| Variable | Grassland specialists |  |  |  |  | Forest specialists |  |  |
| --- | --- | --- | --- | --- | --- | --- | --- | --- |
|  | BOBO | EAME | HOLA | SAVS | KILL | WOTH | EWPE | LEFL |
| *200m* |  |  |  |  |  |  |  |  |
| Water | -0.60 | -1.61 | -0.60 | -0.49 | -2.01 | -1.33 | -2.80 | 4.19 |
| Shrub | -0.92 | 2.22 | -0.65 | -0.64 | -2.50 | 6.85 | 1.75 | -2.87 |
| Wetland | -1.55 | 1.58 | -1.08 | -0.82 | -2.59 | 1.16 | 2.08 | 2.63 |
| Grassland | 46.41 | 40.73 | -2.27 | 34.17 | 3.03 | -2.10 | -0.92 | 4.48 |
| Crops | — | — | — | — | — | -1.60 | -1.12 | -3.89 |
| Cereals | -2.92 | -0.77 | 8.62 | 7.75 | 9.41 | — | — | — |
| Corn | -2.99 | -0.73 | 6.10 | 6.87 | 10.13 | — | — | — |
| Soy | 9.35 | -1.64 | 12.93 | 8.31 | 16.86 | — | — | — |
| Forest | -1.71 | -2.70 | -4.11 | -9.13 | -7.97 | — | — | — |
| Deciduous | — | — | — | — | — | 9.32 | 34.36 | 14.12 |
| Coniferous | — | — | — | — | — | 5.76 | -2.57 | -1.73 |
| Mixed forest | — | — | — | — | — | 16.12 | 6.25 | 4.11 |
| *1km* |  |  |  |  |  |  |  |  |
| Water | -1.21 | -1.95 | -0.99 | -1.15 | -1.41 | -1.04 | -3.56 | -2.46 |
| Urban | -3.33 | -0.82 | -0.80 | -1.08 | -0.95 | -2.00 | -1.60 | -4.03 |
| Shrub | 2.94 | 5.75 | -0.71 | -1.08 | -2.52 | 3.05 | 2.58 | -3.47 |
| Wetland | 2.69 | 6.62 | -0.89 | 1.04 | -2.27 | 4.37 | 6.70 | -1.81 |
| Grassland | 2.33 | 6.72 | 1.24 | 3.24 | 2.70 | -1.84 | 6.35 | -2.37 |
| Crops | — | — | — | — | — | -2.90 | -3.93 | -1.07 |
| Cereals | 4.76 | -3.23 | 10.56 | 1.60 | 8.36 | — | — | — |
| Corn | 1.49 | -3.00 | 9.27 | 2.71 | 5.09 | — | — | — |
| Soy | 1.60 | -3.77 | 11.35 | 5.59 | 3.84 | — | — | — |
| Forest | -2.00 | -4.48 | -1.56 | -1.41 | -1.79 | — | — | — |
| Deciduous | — | — | — | — | — | 12.88 | 2.92 | 2.91 |
| Coniferous | — | — | — | — | — | 1.72 | -5.14 | 5.41 |
| Mixed forest | — | — | — | — | — | 1.07 | -1.69 | 1.61 |
| *Terrain* |  |  |  |  |  |  |  |  |
| Elevation | -0.87 | -5.02 | 13.77 | -2.87 | 4.77 | 4.55 | -3.07 | 5.44 |
| Slope | -1.51 | -1.36 | -1.08 | -1.02 | -1.28 | 0.92 | -2.01 | 8.65 |
| Aspect | 1.32 | 0.83 | 1.43 | 1.39 | 1.92 | 4.64 | 1.59 | -6.55 |
| *Climate* |  |  |  |  |  |  |  |  |
| Bios10 | -3.46 | 3.10 | 10.15 | 2.73 | 5.22 | 11.29 | 4.24 | -10.77 |
| Bios18 | 4.06 | -1.38 | 0.83 | 4.91 | 3.38 | -3.40 | -2.76 | -5.43 |

**Table A6.** Minimum set prioritization results showing the percentage (%) of the distribution and area selected for each feature. Total richness refers to sites within the upper quartile for highest species richness in the focal area.

| Feature | % distribution | area(km<sup>2</sup>) |
| --- | --- | --- |
| Bobolink | 0.30 | 9032 |
| Eastern meadowlark | 0.25 | 6068 |
| Eastern wood Pewee | 0.25 | 22732 |
| Horned lark | 0.25 | 7228 |
| Killdeer | 0.25 | 12040 |
| Least flycatcher | 0.25 | 42992 |
| Savannah sparrow | 0.25 | 11104 |
| Wood thrush | 0.25 | 19032 |
| **Total richness** | 0.25 | 19556 |

**Figure A1.**
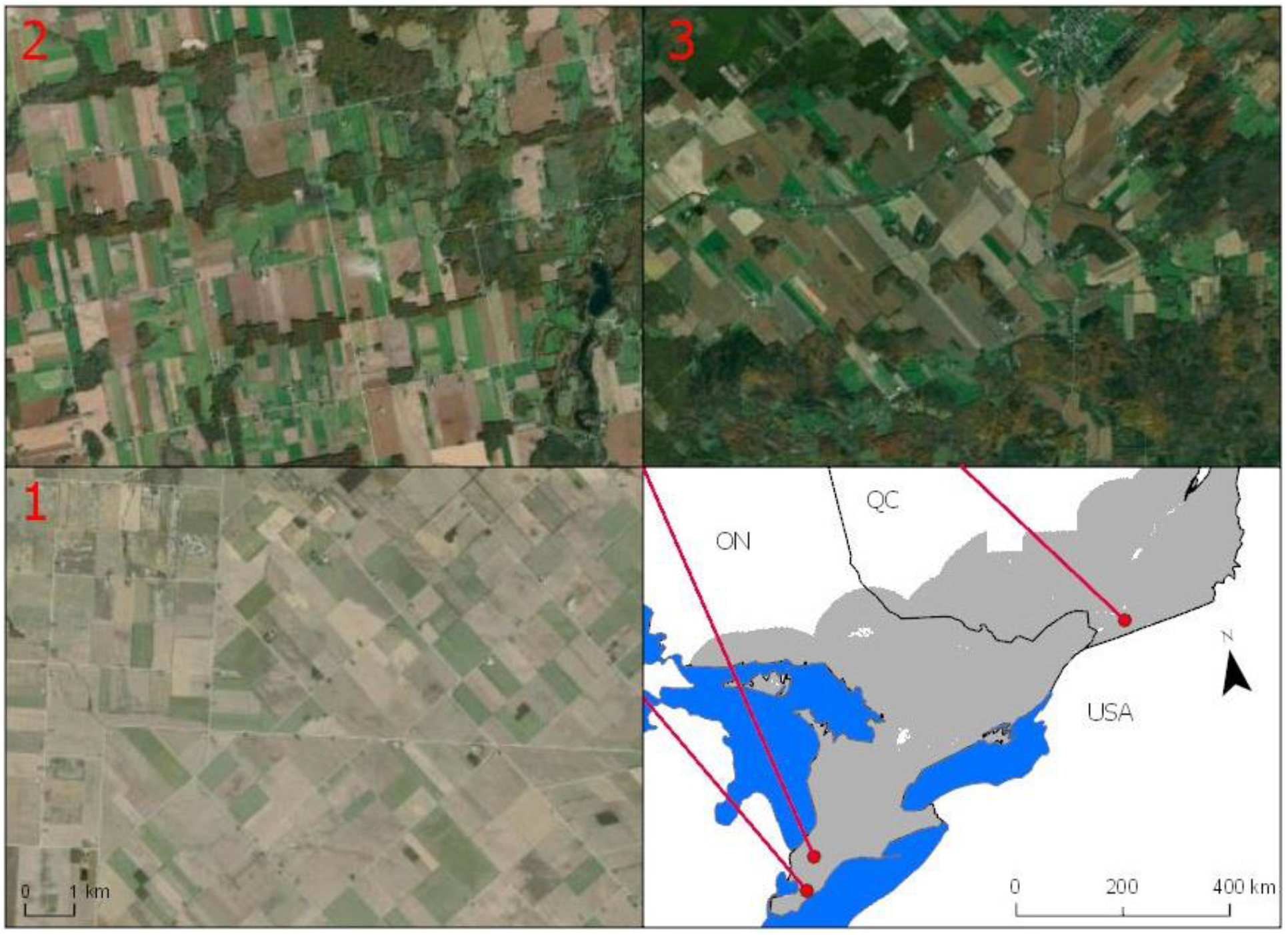
Agricultural landscapes within the study region showing a site with little retained vegetation cover (1) and two sites with vegetation retained in the form of linear woody strips (2) and small forest blocks (3). Site locations are shown in the study region map in the lower right quadrant.

